# The abrogation of condensin function provides independent evidence for defining the self-renewing population of pluripotent stem cells

**DOI:** 10.1101/143339

**Authors:** Alvina G. Lai, Nobuyoshi Kosaka, Prasad Abnave, Sounak Sahu, A. Aziz Aboobaker

**Author notes:** Contributed equally (Alvina G. Lai, Nobuyoshi Kosaka).

## Abstract

Heterogeneity of planarian neoblast stem cells has been categorised on the basis of single cell expression analyses and subsequent experiments to demonstrate lineage relationships. Some data suggest that despite gene expression heterogeneity amongst cells in the cell cycle, in fact only one sub-population, known as *sigma* neoblasts, can self-renew. Without the tools to perform live *in vivo* lineage analysis, we instead took an alternative approach to provide independent evidence for defining the self-renewing stem cell population. We exploited the role of highly conserved condensin proteins to functionally assay neoblast self-renewal properties. Condensins are involved in forming properly condensed chromosomes to allow cell division to proceed during mitosis, and their abrogation inhibits mitosis and can lead to repeated endoreplication of the genome in cells that make repeated attempts to divide. We find that planarians possess only the condensin I complex, and that this is required for normal stem cell function. Abrogation of condensin function led to rapid stem cell depletion accompanied by the appearance of giant cells with increased DNA content. Using previously discovered markers of heterogeneity we show that enlarged cells are always from the *sigma*-class of the neoblast population and we never observe evidence for endoreplication for the other neoblast subclasses. Overall, our data establish that condensins are essential for stem cell maintenance and provide independent evidence that only *sigma*-neoblasts are capable of multiple rounds of cell division and hence self-renewal.

## Introduction

Throughout the lifespan of a multicellular organism with an adult lifespan of months or years, cells in many tissues will turn over, requiring mechanisms in place to control this tissue homeostasis. This requires stem cells of some kind that have the ability to self-renew, proliferate and differentiate into specialised cells for specialised tissue structures. Many animals appear to use germ layer and lineage specified tissue resident stem cells for these processes (Reddien and Alvarado, 2004; Tanaka and Reddien, 2011). Although the potency of such cells in animals varies greatly, most have limited potency. Planarians, however, have at least some individually pluripotent stem cells amongst the broader cycling population of adult stem cells (Wagner et al., 2011), collectively called neoblasts (NBs) (Aboobaker, 2011; Rink, 2013; Ross et al., 2017). NBs give planarians their ability to regenerate any missing tissue structure from small fragments and respond with startling homeostatic plasticity to changing nutritional status (Reddien and Alvarado, 2004; Saló, 2006; González-Estévez et al., 2012a). A current broad definition of NB stem cells is that they represent all cells that are actively cycling, and can be labelled with fluorescent in situ hybridisation (FISH) markers such as *Smedwi-1* (Reddien et al., 2005) and *histone2B (H2B)* (Guo et al., 2006; Solana et al., 2012), which act as pan-NB markers at the transcript level. A number of studies have assayed the genes expressed in NBs and other planarian cells at the whole population level (Blythe et al., 2010; Labbé et al., 2012; Önal et al., 2012; Solana et al., 2012; Kao et al., 2013, 2017) and more recently at the single-cell level (van Wolfswinkel et al., 2014; Issigonis and Newmark, 2015; Wurtzel et al., 2015, 2017; Molinaro and Pearson, 2016; Scimone et al., 2016). This has led to the definition of NB subtypes based on gene expression profiles and revealed that planarians have at least three major subclasses of *smedwi-1+* NBs; with *sigma*-NBs giving rise to both *zeta*-NBs and probably *gamma*-NBs. Each of these is defined by unique expression profiles including enrichment for distinct transcription factors (van Wolfswinkel et al., 2014). In the case of *zinc finger protein 1* (*zfp1*), as well as being a definitive FISH marker of *zeta*-NBs it also reported, through *zfp1(RNAi)* experiments, as being required for transition from *sigma*-NBs to *zeta*-NBs (van Wolfswinkel et al., 2014). Evidence from this study, in particular that genes that define zeta-NB expression are co-expressed in newly minted post-mitotic epidermal progeny and that they increase in expression in NB over the course of S-phase, suggested that *zeta*-NBs may pass through mitosis once to give rise to post-mitotic daughter cells (van Wolfswinkel et al., 2014). This was also supported by the observation that changes in proliferation in response to amputation only impacted sigma-NBs, suggesting that only sigma-NBs, and not other classes of NBs, were self-renewing.

Here, we aimed to provide further independent evidence of *sigma*-NBs being the only NBs capable of multiple rounds of cell divisions and therefore self-renewal. Without the ability to perform transgenesis and *in vivo* lineage tracing to follow stem cell development in planarians, we employed a molecular genetics approach to target proteins that have pivotal roles in regulating cell division that might reveal which cells self-renew and which do not. This led us to study the role of planarian condensins. Condensins are conserved multi-protein complexes essential for chromosome organization, assembly and segregation (Hirano, 2012, 2016). As the mode of action of condensins is specific to M-phase of mitosis or meiosis, perturbation of condensins should only affect cells passing through these cell cycle stages. Two condensin complexes, condensin I and II, have been identified in many animals. Both condensin I and II complexes share the structural maintenance of chromosomes (SMC) 2 and SMC4 subunits (Hirano, 2016) and each complex has their own unique non-SMC subunits: i.e., non-SMC condensin I complex subunit G (NCAPG), non-SMC condensin I complex subunit D2 (NCAPD2) and non-SMC condensin I complex subunit H (NCAPH) for condensin I and similarly NCAPG2, NCAPD3 and NCAPH2 for condensin II (Hirano, 2012). Previous reports have shown that the perturbation of condensins leads to drastic defects in proliferating cells. In plants, abrogation of condensins resulted in enlargement of endosperm nuclei (Liu et al., 2002). Loss of condensin I in zebrafish causes chromatid segregation defects in neural retina progenitors and polyploidization (Seipold et al., 2009). In mammalian embryonic stem (ES) cells, simultaneous depletion of both condensin I and II resulted in the accumulation of enlarged interphase nuclei (Fazzio and Panning, 2010). Although complete elimination of condensin proteins appear to have lethal effects, reduction in condensin levels may be tolerated to allow functional studies on surviving cells (Gosling et al., 2007; Longworth et al., 2008; Murakami-Tonami et al., 2014; Nishide and Hirano, 2014; Frosi and Haering, 2015). The function of condensins in planarian neoblasts is not known. In light of this, we sought to confirm the contribution of condensins to NB proliferation. We reasoned that condensin depletion may lead to endoreplication of cells that remain in the cell cycle and self-renew. Cell that do not pass through mitosis but leave the cell cycle and differentiate would be depleted, as their own populations are not renewed by differentiation of daughter progeny from self-renewing stem cells.

We demonstrate that condensin I genes in the model planarian *Schmidtea mediterranea* have enriched expression in stem cells and are essential for tissue homeostasis and regeneration. RNA interference (RNAi)-mediated knockdown of all five condensin subunits resulted in a drastic decline in NBs. Remaining NBs positive for the stem cell marker *smedwi-1* or *H2B* in RNAi animals are often morphologically enlarged and have increased DNA content. These enlarged NBs are only ever positive for the *sigma*-class NB marker and never the *zeta*- or *gamma*-class markers. Enlarged *sigma*-NBs have increased DNA content but are non-mitotic, indicating that these cells may have undergone endocycling as a result of condensin depletion. Our results provide independent evidence that *sigma*-NBs are the only population of stem cells capable of multiple rounds of cell division and hence self-renewal in *S. mediterranea*.

## Materials and methods

### Phylogenetic analyses

*S. mediterranea* condensin orthologs were identified by tBlastn against the planarian transcriptome and genome (Robb et al., 2008, 2015) using condensin protein sequences from *Drosophila melanogaster, Homo sapiens* and *Mus musculus* as queries. Condensins from other flatworm species *(Schmidtea polychroa, Planaria torva, Dendrocoelum lacteum, Polycelis nigra* and *Polycelis tenuis*) were obtained from Planmine (Brandl et al., 2016). Multiple sequence alignments of condensin proteins were performed using MAFFT (Katoh et al., 2009) followed by phylogenetic tree generation using RAxML (Stamatakis, 2014) using the gamma WAG model with 1000 bootstrap replicates. The best-scoring maximum likelihood tree was generated. The tree figure was made using Geneious (Kearse et al., 2012). Complete list of transcript sequences obtained are listed in Dataset S1.

### Cloning of condensin family genes

*S. mediterranea* condensin family genes identified above were cloned into the double-stranded RNA expression vector (pT4P) as previously described (Rink et al., 2009). Colony PCR was performed using the M13 forward and reverse primers followed by Sanger sequencing using the AA18 or PR244 primer. Complete list of primer sequences used for PCR and cloning are listed in Table S1.

### Animal culture

Asexual *S. mediterranea* animals were cultured at 20°C in 1X Montjuic salts (Cebrià and Newmark, 2005). The 1X Montjuic salt solution was prepared using milliQ ddH2O with the following composition: 1.6mM NaCl, 1mM CaCl_2_, 1mM MgSO_4_, 0.1mM MgCl_2_, 0.1mM KCl, 1.2mM NaHCO_3_. The worms were fed with organic beef liver once a week and were starved for 1 week prior to any experimental procedures to minimize non-specific background from gut contents. Animals were kept in the dark at all times apart from during feeding and water changing.

### RNA interference (RNAi) by injection

All RNAi experiments were performed on 3 to 4mm worms, according to procedures previously described (Felix and Aboobaker, 2010). Control RNAi was performed by injecting worms with double stranded (ds) RNA encoding for green fluorescent protein (GFP) that is not present in *S. mediterranea*. Condensin dsRNA was prepared by *in vitro* transcription of the gene amplified from the pT4P vector mentioned above with primers listed in Table S1. The GFP gene was amplified from the pGEMT-GFP plasmid for use as control during RNAi (Solana et al., 2012). PCR products were purified using the Wizard SV Gel and PCR clean-up kit (Promega). This was then followed by in vitro transcription using T7 RNA polymerase (Roche). The in vitro transcribed dsRNAs were treated with Turbo DNAse (Ambion), lithium chloride precipitated in the presence of glycogen and re-suspended in molecular grade water to a final concentration of 2mg/mL. 1 week starved animals were injected using the Nanoject II microinjector (Drummond Scientific, Oxford, UK) for three consecutive days in the first week and another three consecutive days in the second week. Worms were monitored throughout the procedure and day 1 post RNAi in all experiments is considered to be the first day after the sixth dsRNA injection. All experiments were performed in triplicates and at least 10 worms were used for each time point.

### Gamma irradiation

Gamma irradiation was performed according to previously described procedures (González-Estévez et al., 2004). Asexual *S. mediterranea* animals were irradiated at 30Gy in a sealed ^137^Cs source. All experiments were performed in triplicates and at least 10 worms were used for each time point.

### Fluorescent in situ hybridization (FISH)

Whole-mount FISH (WISH) and double FISH (dFISH) were performed according to previously described procedures (King and Newmark, 2013) with the following modifications. For sigma-, zeta- and gamma-classes dFISH, N-acetyl cysteine (NAC) treatments on worms were done for 10 minutes. For proteinase K treatment, animals were for 15 minutes followed by 30 minutes of post-fixation with 4% formaldehyde. Saline sodium citrate (SSC) washes were performed as follows: two times washes with 2X SSC, two times washes with 0.2X SSC and two times washes with 0.05X SSC (each wash was for 20 minutes). Fast blue-based detections were performed as described in previous work (Lauter et al., 2011; Cowles et al., 2013). Detection of sigma-, zeta- and gamma-classes were performed with pooled probes: sigma-class *(soxP-1* and *soxP-2*), zeta-class (*zfp-1 and soxP-3*) and gamma-class (*gata4/5/6* and *hnf-4*). Primer sequences for NB subclasses and NB progenitor FISH detection were obtained from previous reports (Solana et al., 2012; van Wolfswinkel et al., 2014). All experiments were performed in triplicates and at least 10 worms were used for each time point.

### Immunohistochemistry and TUNEL assay

Immunolabeling of phospho-histone-3 (H3P) was performed according to a previously described protocol with some modifications (Newmark and Alvarado, 2000; González-Estévez et al., 2012b). Worms were killed in 2% hydrochloric acid/Holtfreter’s solution followed by fixation in Carnoy’s solution. Primary antibody used was rabbit anti-phospho-histone-3 (Millipore) at a 1:1000 dilution. Anti-rabbit HRP (Abcam) secondary antibody (1:4000 dilution) was used followed by tyramide signal amplification according to a previous protocol (King and Newmark, 2013). TUNEL was performed as previously described (Pellettieri et al., 2010) with modifications described in other studies (Almuedo-Castillo et al., 2014; Tejada-Romero et al., 2015). All experiments were performed in triplicates and at least 10 worms were used for each time point.

### Image analyses

Bright field images were captured using the Zeiss SteREO Discovery V8 microscope attached with a Canon EOS 1200D digital SLR camera. Fluorescent images were taken with an inverted Olympus FV1000 confocal microscope and the Zeiss 880 Airyscan microscope. Bright field images were processed with the Adobe Photoshop CS6 software (with image backgrounds corrected to black or white) and confocal stacks were processed using the Fiji processing package. Image compositions were done with the Adobe Illustrator CS5 software. Cell counts were performed on confocal Z stacks at 40X magnification from dorsal to ventral, normalised by the area measured based on OIB files metadata. Measurements of cell areas in μm^2^ were performed in Fiji (Schindelin et al., 2012). Graphs and statistical test results were generated using R ggplot2 (Wickham, 2009).

### Cell dissociation and fluorescence activated cell sorting (FACS)

Cell dissociation was performed as previously described (Romero et al., 2012) with the following modifications. For DNA staining, Hoechst 34580 (Sigma) was used (1mg/mL stock concentration) at a final concentration of 20uL/mL. Cytoplasmic staining was performed using calcein AM (Thermo Fisher Scientific; 1mg/mL stock concentration) at a final concentration of 0.5uL/mL. Staining was performed for 2 hours at room temperature. Just prior to FACS acquisition, 1uL/mL of propidium iodide (Sigma) solution was added to the cells. FACS was performed with the BD Aria III instrument. Hoechst 34580 signal was collected at 450nm while calcein AM signal was collected at 488nm.

## Results and Discussion

### Condensin I subunits are required for tissue homeostasis and regeneration in planarians

To ascertain the biological roles of condensins in NBs, we set out to perform an initial survey of condensin subunit genes in planarians using extant genomic and transcriptomic resources (Robb et al., 2008, 2015, Kao et al., 2013, 2017; Brandl et al., 2016). From our search, we were able to only identify orthologs of condensin I subunits *(SMC2, SMC4, NCAPG, NCAPD2* and *NCAPH*; Fig. 1A, B). Similarly, in other triclad flatworms *(Dendrocoelum lacteum, Planaria torva, Polycelis tenuis, Polycelis nigra* and *Schmidtea polychroa*), we only identified protein subunits of condensin I (Dataset S1). Thus, it appears that triclad flatworms do not have orthologs of condensin II subunits, which are present in other lophotrochozoans. Both *Drosophila* and *Caenorhabditis* have both condensin I and II complexes. It is thought that the last common ancestor of eukaryotes possessed both condensin I and II, and condensin II has been independently loss multiple times during evolution (Hirano, 2016).

**Fig. 1.**
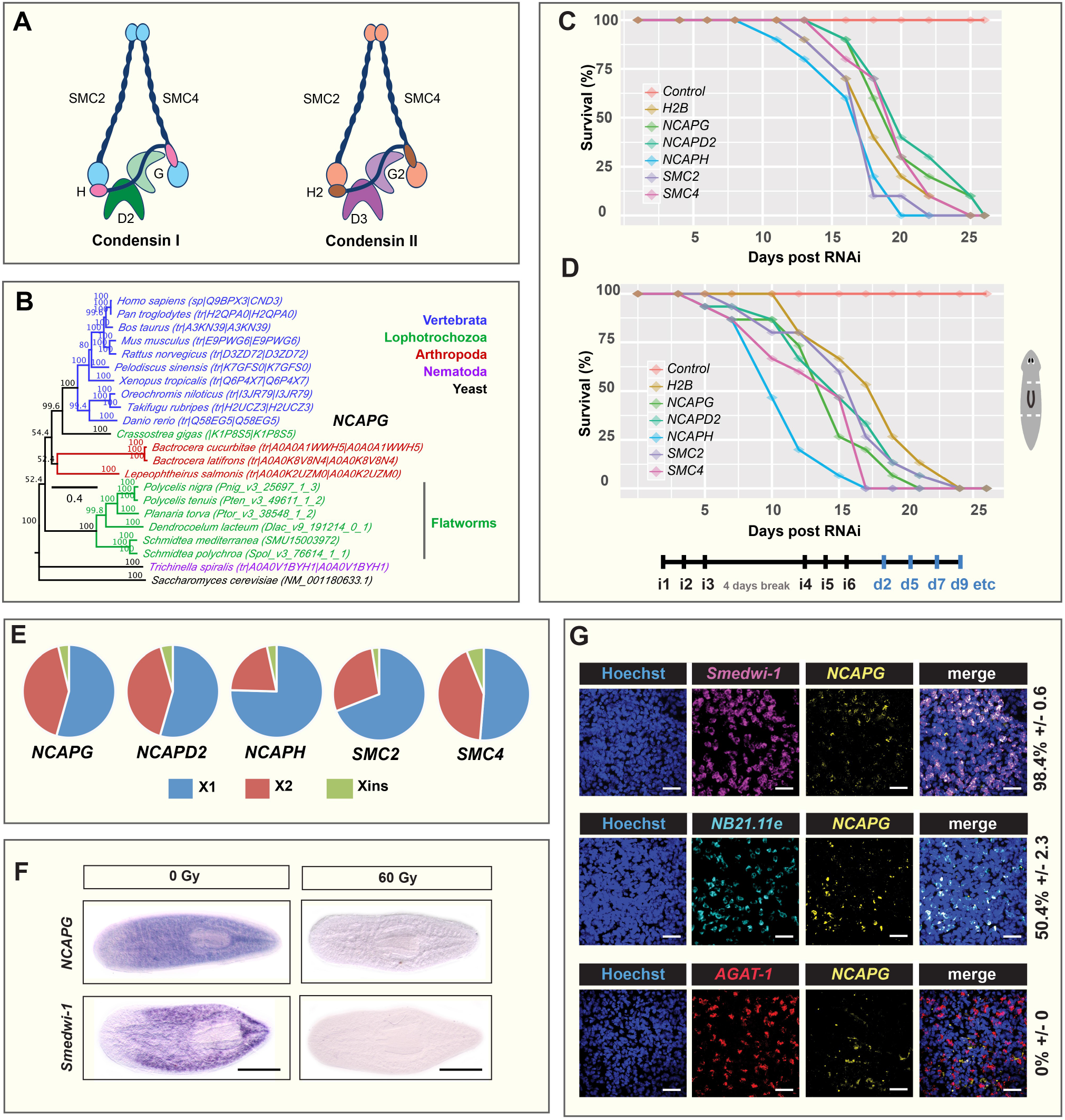
Planarian condensin I subunits are expressed in NBs and are required for regeneration and tissue homeostasis. **(A)** Subunit composition of condensin I and condensin II. Schematic diagram adapted from (Hirano, 2016). **(B)** Phylogenetic tree of *Smed-NCAPG* and orthologs from representative species. Survival graphs of condensin I subunits RNAi animals under **(C)** homeostatic and **(D)** regenerative conditions. Experimental timeline indicating injection schedule denoted as iX and days post injection denoted as dX. RNAi worms were amputated pre-and post-pharyngeally. **(E)** Expression proportion values of condensin I subunits in *S. mediterranea* X1, X2 and Xins cell populations based on meta-analyses of multiple RNA-sequencing datasets (Kao et al, 2017). **(F)** WISH analysis of *NCAPG* two days after 60 Gray (Gy) irradiation. The *smedwi-1* marker was used to probe for the presence of NBs with (60 Gy) and without (0 Gy) irradiation. **(G)** DFISH was performed to ascertain *NCAPG* expression patterns in NBs and post-mitotic progeny. Numbers indicate the percentage of NBs, early progeny (*NB21.11e*) or late progeny (*AGAT-1*) coexpressing *NCAPG* (N=300 cells per dFISH ± standard error). Scale bars represent 50μm.

We performed independent RNAi-mediated knockdown of all five condensin I subunits in *S. mediterranea*. We observed lethality during homeostasis (Fig. 1C) and amputated fragments that failed to regenerate (Fig. 1D). RNAi worms exhibited head regression during homeostasis, dorsal lesioning and the inability to form blastemas during regeneration (Fig. S1A,B). Knock-down of condensin I subunits phenocopied previous observations of *Smed-H2B(RNAi)* animals suggesting that a rapid and potent NB impairment has taken place in each case (Solana et al., 2012) (Fig. S1A,B).

### Condensin I subunits are expressed in planarian stem cells

Phenotypes of condensin I RNAi animals hinted at a broad NB defect. Reports in other systems suggest that ES cells and somatic cells have distinct dependence on condensin complex activity. ES cells, but not somatic cells are highly susceptible to reduced levels of condensins (Fazzio et al., 2008; Hu et al., 2009; Fazzio and Panning, 2010) perhaps because they have dynamic chromatin with only loosely bound architectural proteins (Meshorer and Misteli, 2006). This prompted us to investigate the expression profiles of condensin I subunits across different planarian cell populations. We first utilised expression proportion values obtained from a meta-analysis performed using RNA-sequencing datasets from multiple laboratories (Kao et al., 2017). Planarian cells are broadly categorised into three subpopulations: X1 representing S/G2/M NBs, X2 representing G1 NBs and post-mitotic progeny and Xins representing irradiation-insensitive post-mitotic differentiated cells (Hayashi et al., 2006). We observed that all five condensin I subunits had the majority of transcript expressed in the X1 NBs: *NCAPG* (54%), *NCAPD2* (55%), *NCAPH* (79%), *SMC2* (69%) and *SMC4* (51%; Fig. 1E). Whole mount in situ hybridization (WISH) of *Smed-NCAPG* on wild-type animals revealed a NB-like expression pattern and following gamma irradiation, we observed a loss of *Smed-NCAPG* as a consequence of NB elimination (Fig. 1F). We found that over 90% of cells expressing *Smedwi-1* co-express transcripts from all five condensin 1 complex genes, while between 23 to 69% of cells expressing *NB-21.11e* (an early marker of progeny committed to epidermal cell fate) co-express condensins (Fig. 1G, Fig. S1C, D, E, F). However, cells expressing the later epidermal fated progeny marker *AGAT-1* are all negative for condensin subunit expression (Fig. 1G, Fig. S1 C, D, E, F). The observed expression in *NB-21.11e* cells suggests that residual condensin transcription is still detectable upon mitotic exit but completely disappears as cells differentiate further to *AGAT-1* expressing cells. Overall, our results suggest that condensins are mostly expressed in proliferating cells, in line with their mode of action in regulating cell division, but transcripts persists after cell cycle exit.

### Knockdown of *Smed-NCAPG* results in abnormal karyotype and depletion of mitotic stem cells

In other eukaryotic systems, loss of any condensin subunits causes segregation and chromosome condensation defects (Saka et al., 1994; Strunnikov et al., 1995; Ouspenski et al., 2000; Lavoie et al., 2002; Lam et al., 2006; Seipold et al., 2009), thus, it seems likely that RNAi animals would harbour these defects. From our karyotyping analyses, we observed aberrant chromosomal morphologies in *Smed-NCAPG(RNAi)* animals e.g. chromosome fusions (Fig. 2A), decondensed chromosomes and elongated chromatids and abnormal numbers of chromosomes, probably as a result of gross nondisjunction events (Fig. S1G). Given the high proportion of *Smed-NCAPG* expression in X1 NBs and the lethality of *Smed-NCAPG (RNAi)* animals, we sought to determine the extent and nature of NB perturbation in these animals. We performed fluorescence-activated cell sorting (FACS) analyses on *Smed-NCAPG (RNAi)* animals and observed a time-dependent decrease in X1 and X2 cells over three post-RNAi time points (Fig. 2B). Xins cells appeared to be unaffected by *Smed-NCAPG (RNAi)*. Thus, depletion of *Smed-NCAPG* affects dividing cells as predicted, but also results in a reduction of transient post-mitotic progeny. As independent confirmation, we performed WISH using the pan-neoblast markers *Smedwi-1+* (Reddien et al., 2005) and *H2B+* (Solana et al., 2012) in *Smed-NCAPG(RNAi)* animals and we observed, in line with FACS results, NBs are lost by 9 days after RNAi (Fig. 2C, D, E). These observations coincided with a decline in mitotic stem cells detected by phosphorylated histone H3 (H3P) immunolabeling. (Fig. 2F, G). We also assessed the levels of apoptosis using the terminal deoxynucleotidyl transferate dUTP nick end labelling (TUNEL) assay to see if our results where indicative of raised levels of NB apoptosis. At day 1 post-RNAi, *Smed-NCAPG(RNAi)* worms already had increased levels of cell death correlating with NB depletion (Fig. 2H). Together these data confirmed a defects maintenance and survival of NBs. From these data we conclude that NB depletion may be caused by aberrant mitotic events leading to cell death or by stem cells differentiating at raised levels without self-renewal or some combination of the two. From our WISH experiment of *NB21.11e* and *AGAT-1* in *Smed-NCAPG(RNAi)* worms in days 2, 5 and 9 post-RNAi, we observed a time-dependent depletion of progenitor cells, suggesting that depletion is concomitant with NB lost indicating that NB depletion rather than increased differentiation might be responsible (Fig. S2 A, B, C, D).

**Fig. 2.**
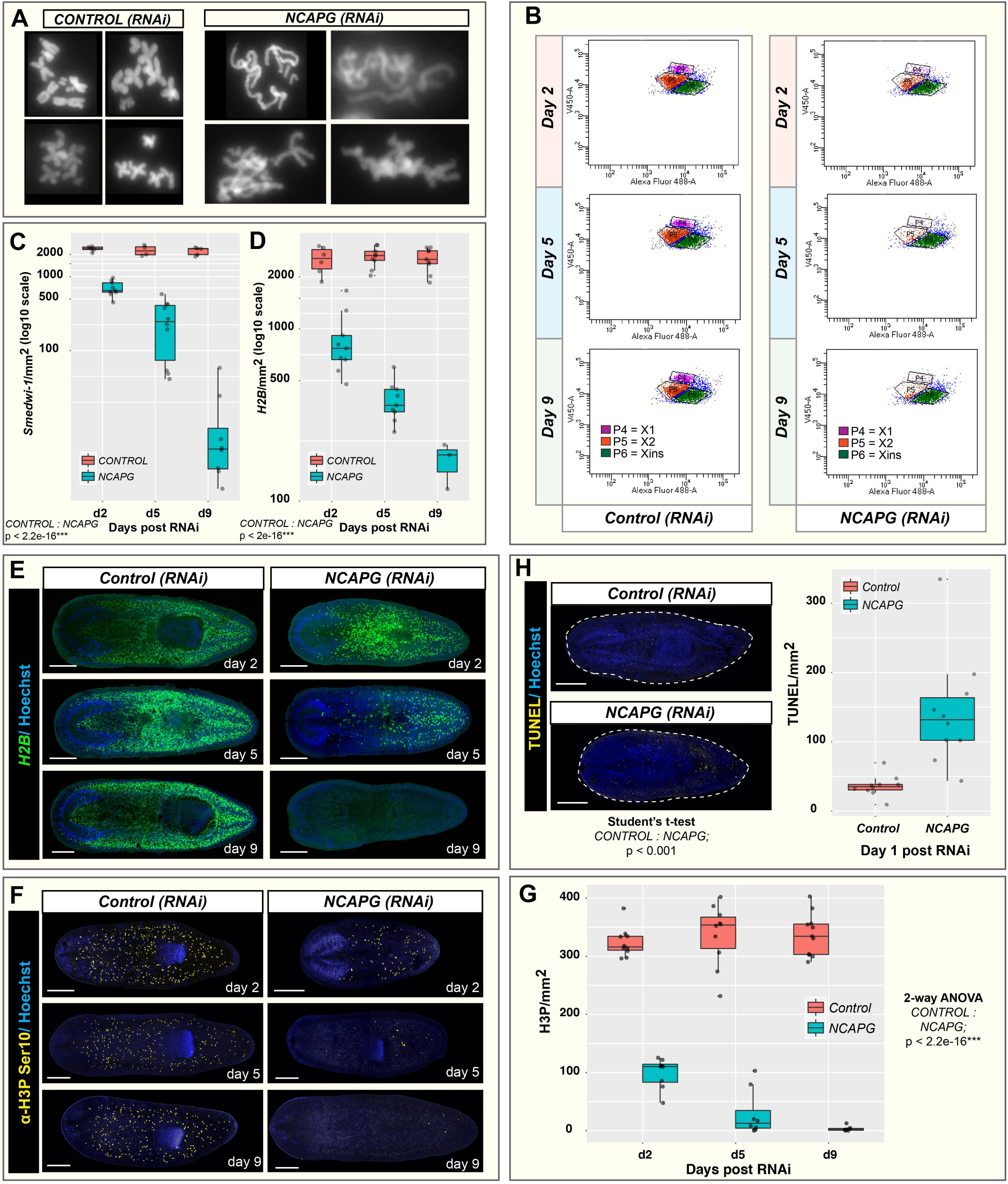
*Smed-NCAPG (RNAi)* resulted in gross chromosomal defects and is required for NB homeostasis. **(A)** Karyotype of *Smed-NCAPG (RNAi)* animals showing chromosomal fusion events. **(B)** FACS plots of Hoechst area (DNA stain) versus Calcein area (cytoplasmic stain) allow the identification of X1 (purple), X2 (orange) and Xins (green) populations. Depletion of *NCAPG* resulted in a time-dependent decrease in X1 and X2 cells. Quantification of **(C)** *Smedwi-1+* and **(D)** *H2B+* stem cells in homeostatic animals at 2, 5 and 9 days post RNAi. **(E)** WISH was performed on homeostatic *Smed-NCAPG (RNAi)* animals using the stem cell marker *H2B*. Mitosis in homeostatic *Smed-NCAPG (RNAi)* animals were detected using **(F)** H3P immunolabeling techniques and **(G)** cell numbers were quantified at selected time points post RNAi. **(H)** Cell death *Smed-NCAPG (RNAi)* animals were detected using TUNEL and cell numbers were quantified at 1 day post RNAi. Scale bars represent 0.5mm. *** p<0.001 (2-way ANOVA).

The depletion of *Smedwi-1+* and *H2B+* NBs in *Smed-NCAPG (RNAi)* animals could either reflect a global decrease in all NBs or specific NB subclasses (van Wolfswinkel et al., 2014). Multiple lines of evidence suggest that *zeta*- and *gamma*-NBs are derived from *sigma*-neoblasts. Grafts from *zfp-1(RNAi)* donors transplanted into NB-depleted hosts could regenerate the *zeta-* class from *sigma*-NBs (van Wolfswinkel et al., 2014). Since sigma-NBs are the source of the zeta-class NBs and the likely source of gamma-class NBs, depletion of the former would also affect the latter. We performed double fluorescent WISH (dFISH) of *sigma (soxP-1* and *soxP-2*), *zeta* (*zfp-1 and soxP-3*) and *gamma (gata4/5/6* and *hnf-4*) with *Smedwi-1* in *Smed-NCAPG (RNAi)* animals and saw marked decreases in all three NB subclasses (Fig. 3, Fig. S2 E, F). FISH measuring depletion of sigma-NBs in RNAi worms of the four other condensin genes produced similar results, with drastic depletion of sigma-NBs (Fig. S3A, B). Overall these data that RNAi based inhibition of condensing leads to rapid depletion of NBs and is accompanied by defects in chromosome condensation and mitosis.

**Fig. 3.**
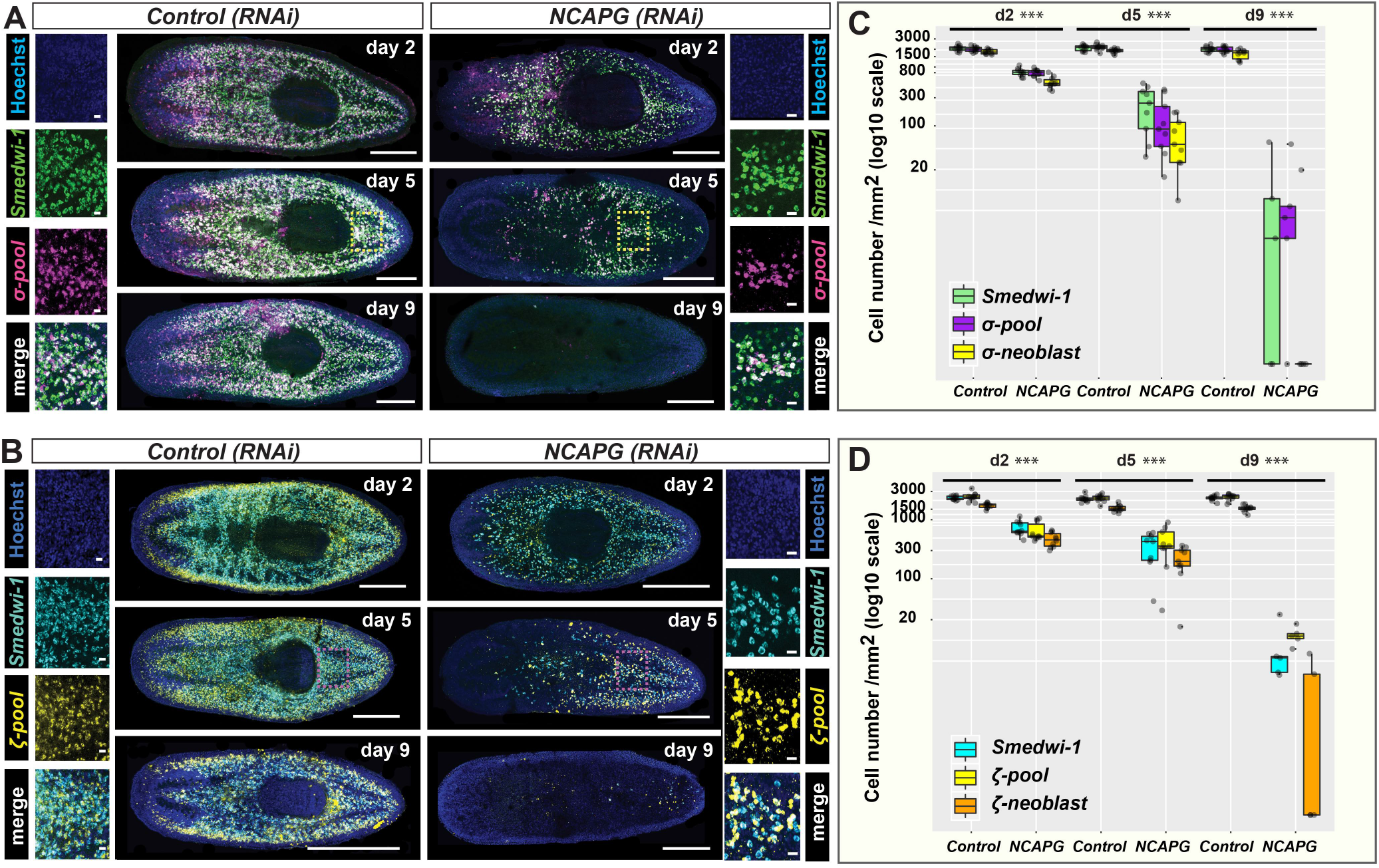
*Smed-NCAPG (RNAi)* results in the depletion of sigma-and zeta-NBs. Lineage markers labelling **(A)** sigma-neoblasts and **(B)** zeta-neoblasts were assessed by dFISH in RNAi animals at 2, 5 and 9 days post RNAi. The day-9 time point depicts maximal perturbation of sigma-and zetaneoblasts. Yellow and pink boxes indicate the regions representing the zoomed images. Scale bars in the whole mount images represent 0.5mm. Scale bars in the zoomed images represent 20μm. Quantification of **(C)** sigma-neoblasts and **(D)** zeta-neoblasts at 2, 5 and 9 days post RNAi. ***p < 0.001 (Student’ s t-test; pairwise comparisons between each cell population in control versus RNAi animals).

### Knockdown of *Smed-NCAPG* results in the endocycling of stem cells

From our phenotypic assessment of any remaining NBs in RNAi animals at 5 days post injection, we observed many enlarged *H2B+* cells (Fig. 4A, D). Although these enlarged NBs have increased areas of DNA staining and appear to be multinucleated, they were always negative for H3P and thus are not mitotic, perhaps never successfully entering M-phase due abrogation of condensin function (Fig. 4A). We only ever observed H3P+ signal in normal sized NBs in *Smed-NCAPG (RNAi)* animals (Fig. 4A, E), suggesting some NBs can enter M-phase. This suggested that endoreplication has occurred in some NBs upon depletion of *Smed-NCAPG*. Using markers of NB sub-classes we found that enlarged stem cells co-express *Smedwi-1* and the sigma-class markers. (Fig. 4C). Cell area measurements revealed that enlarged NBs are only positive for the sigma-class markers, but not the zeta- or gamma-class markers (Fig. 4F,G, Fig. S2G). Analyses on *Smed-NCAPD2, NCAPH, SMC2* and *SMC4* (RNAi) animals also revealed that enlarged NBs are in fact sigma-NBs (Fig. S3C, D). A variant of endoreplication known as endocycling happens when cells undergo successive rounds of genomic DNA replication without mitosis or nuclear envelope breakdown, chromosome condensation or cytokinesis and results in somatic polyploidy (Edgar et al., 2014). Indeed, as we observed that enlarged sigma-NBs in *Smed-NCAPG(RNAi)* animals are also non-mitotic (Fig. 4B), it seems likely enlarged cells have undergone endocycling after abrogation of condensin function.

**Fig. 4.**
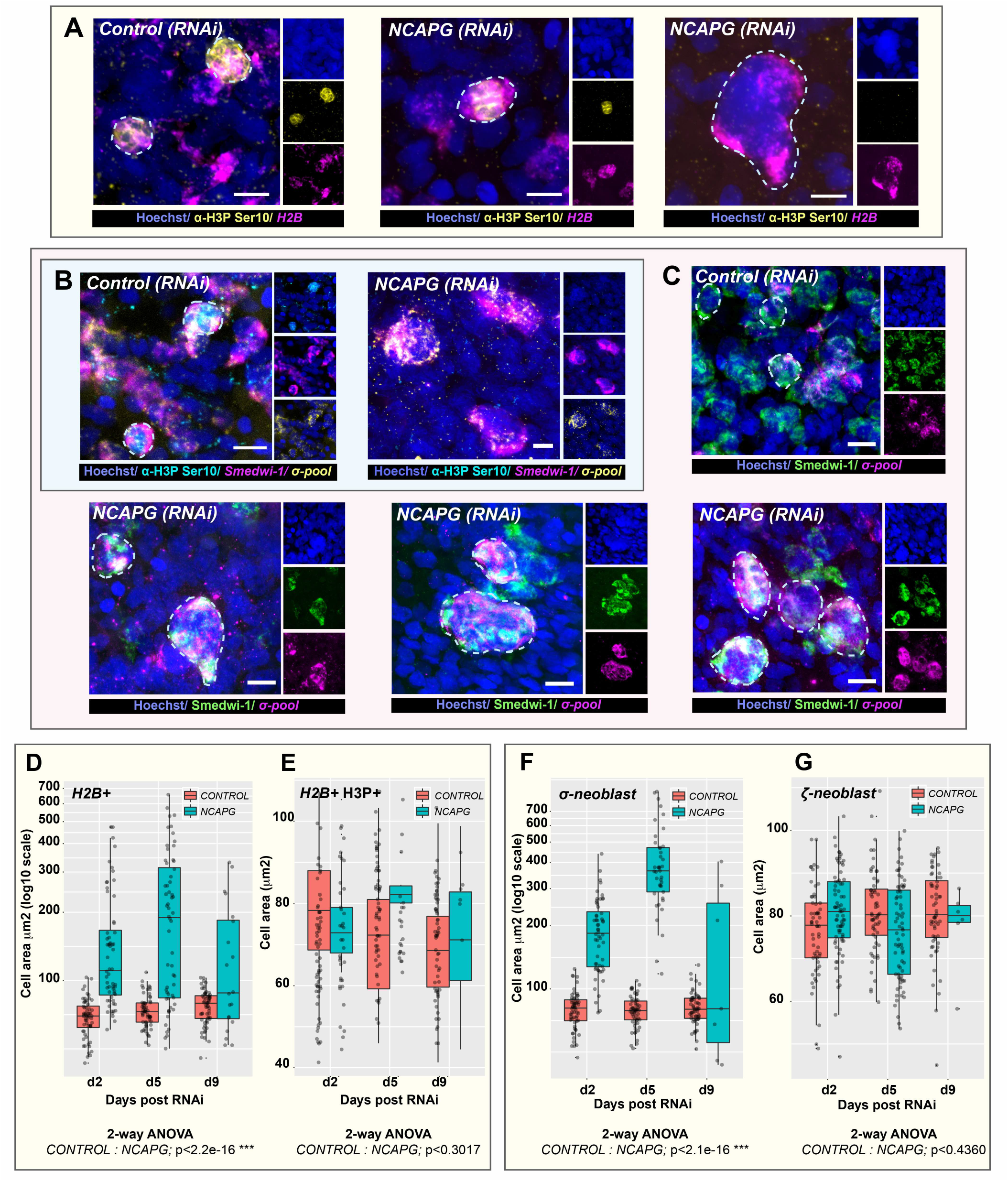
*Smed-NCAPG (RNAi)* results in the formation of enlarged sigma-NBs. **(A)** Depletion of *NCAPG* resulted in enlarged stem cells in the absence of mitosis. Enlarged stem cells are always negative for the H3P mark (far right image) while a subset of morphologically normal stem cells could still undergo mitosis in RNAi animals (middle image). **(B)** Triple labelling of *Smedwi+, sigma+* and H3P+ cells. *Smed-NCAPG (RNAi)* animals show enlarged sigma-neoblasts that are non-mitotic. **(C)** A subset of sigma-neoblasts are enlarged and multinucleated in RNAi animals. Scale bars represent 10μm. Quantification of **(D)** H2B+, **(E)** H2B+/H3P+, **(F)** sigma-neoblasts and **(G)** zeta-neoblasts cell sizes in RNAi versus control animals at 2, 5 and 9 days post RNAi. *** p<0.001 (2way ANOVA).

The observation of multi-nucleated enlarged sigma-NBs, but not zeta- or gamma-NBs, provides independent evidence that this population is the only NB subclass that can undergo multiple rounds of cell division and therefore self-renewal. After condensin complex abrogation in RNAi animals some *sigma*-NBs, that do not die after mitosis fails, continue to attempt to replicate and self-renew, accumulate replicated DNA. We never observe enlarged zeta- or gamma-NBs suggesting they do not go through multiple rounds of division and are post-mitotic early after a single division. Indeed zeta-NB markers are also co-expressed with post-mitotic progeny in single cell expression assays and *zfp1* at least is co-expressed with other early progeny markers (van Wolfswinkel et al., 2014; Abnave et al., 2016; Molinaro and Pearson, 2016). One potential possibility that remained extant from previous work is that zeta-NB or gamma-NB markers switch on and off in S and G1 phases of the cell cycle respectively. So although are expressed mainly in cells that will imminently leave the cell cycle, they also transiently expressed in some portion of repeatedly cycling NBs. Our work here provides strong further evidence against this as enlarged zeta or gamma-NBs are never observed. Instead a subset of sigma-NBs during S phase gain either zeta or gamma expression signatures, proceed through M-phase and then give rise to zeta- and gamma class (non-NB) cells which are post-mitotic (van Wolfswinkel et al., 2014). Future work will clearly be directed at understanding the regulatory mechanisms that prompt sigma-NBs to make these decisions as well as uncovering all the different decisions made for different lineages.

## Conclusion

This study represents the first functional analysis of condensin I subunits in adult stem cells in an in vivo model system. We show that triclad planarians have a single condensin I complex. *S. mediterranea* condensins are expressed in stem cells and are essential for stem cell homeostasis and tissue regeneration (Fig. 1). Upon depletion of condensin genes, we observed gross chromosomal defects, increased stem cell death and the appearance of surviving stem cells with abnormal morphology (Fig. 2, 3). Knockdown of condensins affects the ability of cells to successfully undergo M-phase of the cell cycle as chromosome condensation fails. In planarians we observe large cells with increased DNA content and the continued absence of H3P staining (Fig. 2A and Fig. 4A, B). This phenotype, while predictable based on previous studies of condensins, allowed us to gain independent evidence for the nature of different NB sub-classes previously defined by expression patterns and evidence of lineage relatedness from functional experiments (van Wolfswinkel et al., 2014). We have shown that sigma-NBs, but not zeta- or gamma-NBs, are the only NBs currently defined by differing expression profiles that undergo multiple rounds of endocycling after condensin depletion, suggesting that they are also the only NBs that can undergo multiple rounds of cell division and hence self-renewal (Fig. 4, Fig. S3C, D). These data show therefore that, so far, there are no markers of stem cell heterogeneity that demonstrate differences amongst cells that undergo more than one pass through M-phase of the cell cycle in planarians, and current NB-subclasses consist of self-renewing potentially pluripotent cells (*sigma*-NBs) and lineage committed NBs, which themselves likely only divide once to produce post-mitotic lineage specified daughter progeny.

## Author contributions

A.A.A., A.G.L. and N.K. conceptualized and designed the study to assess self-renewal. A.G.L., N.K., P.A. and S.S. performed the experiments. A.G.L. and N.K. analysed the data. A.A.A., A.G.L. and N.K. wrote the initial manuscript draft. A.G.L. and A.A.A. finalised the submitted draft. All authors were involved in reviewing and editing the final manuscript.

## Acknowledgements

We thank Aboobaker lab members for advice and comments on the manuscript.

## Supplementary figure legend

**Fig. S1.**
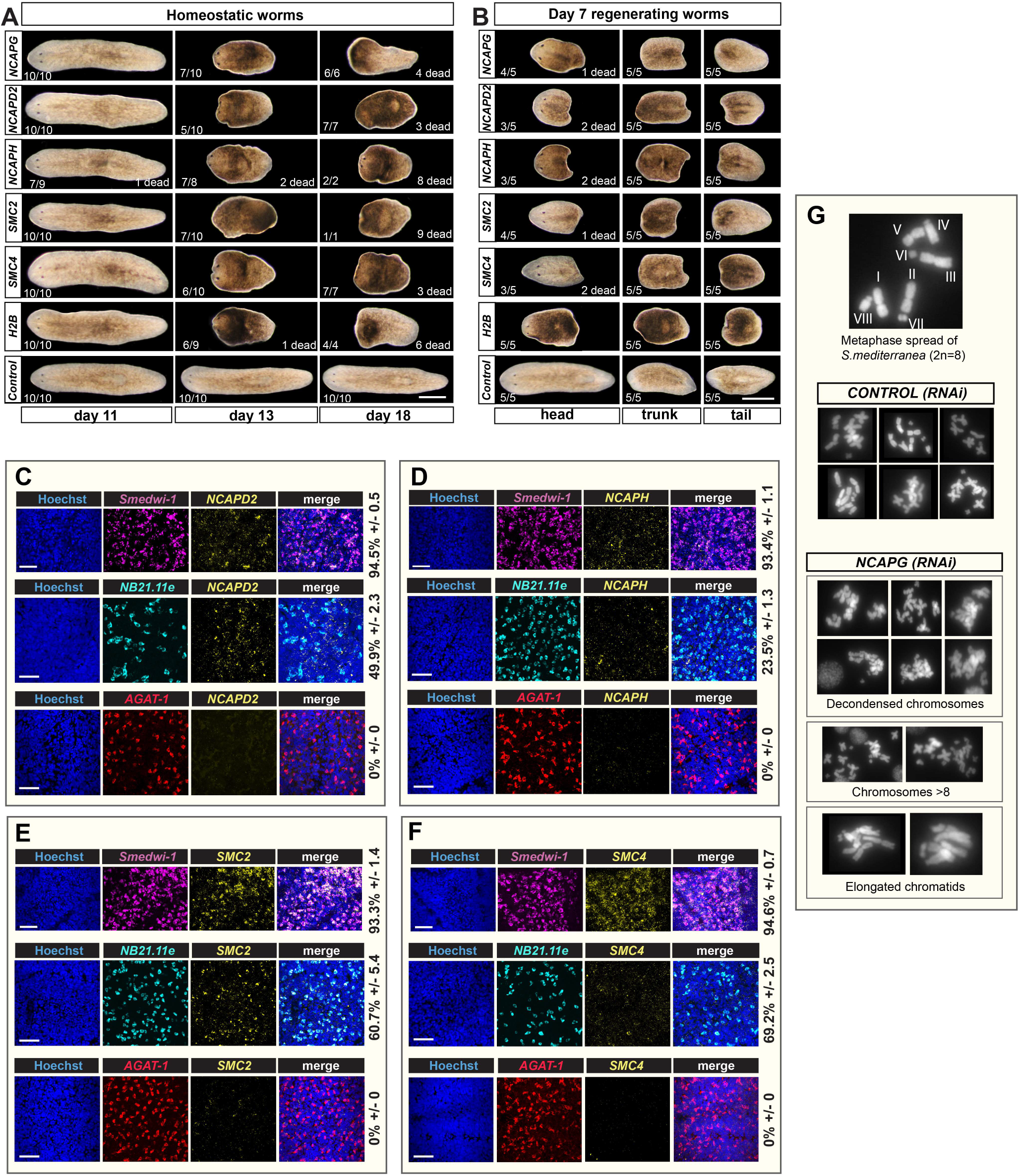
RNAi of condensin I subunits results in impaired tissue homeostasis and regenerative ability. RNAi-mediated knock down of condensin I subunits resulted in animals with **(A)** impaired tissue homeostasis and **(B)** regenerative capacity. RNAi animals failed to form any blastema, were unable to regenerate and they eventually die. Scale bars represent 0.5mm. Expression patterns of **(C)** *NCAPD2*, **(D)** *NCAPH*, **(E)** *SMC2* and **(F)** *SMC4* in stem cells and post-mitotic progenies were determined using dFISH. Numbers indicate the percentage of stem cells, early progeny (*NB21.11e*) or late progeny (*AGAT-1*) co-expressing each respective condensin subunit gene (N=300 cells per dFISH ± standard error). Scale bars represent 50μm. **(G)** Additional chromosomal abnormalities resulted from the depletion of *NCAPG*. The top image represents an example of metaphase spread from wild-type *S. mediterranea* cells.

**Fig. S2.**
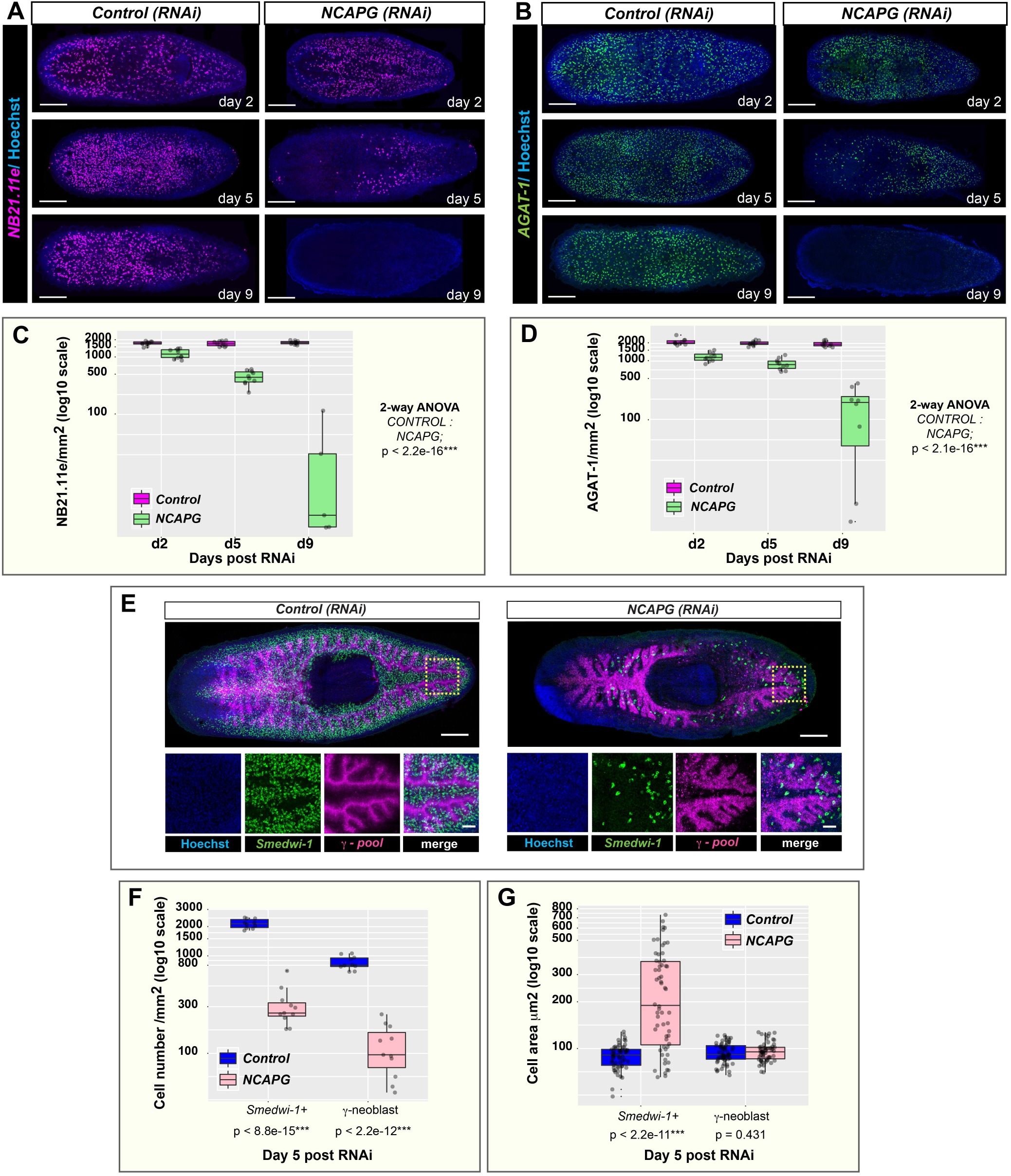
*Smed-NCAPG (RNAi)* results in the depletion of early and late progeny and gamma-neoblasts. WISH was performed on homeostatic *Smed-NCAPG (RNAi)* animals using markers for **(A)** *NB21.11e* cells and **(B)** *AGAT-1* cells at 2, 5 and 9 days post RNAi showing a time-dependent depletion of both cell populations. Scale bars represent 0.5mm. Quantification of **(C)** *NB21.11e* cell numbers and **(D)** *AGAT-1* cell numbers taken at the same time points. Lineage marker labelling **(E)** gamma-neoblasts was assessed by dFISH in RNAi animals at 5 days post RNAi. Yellow boxes indicate the regions representing the zoomed images. Scale bars in the whole mount images represent 0.5mm. Scale bars in the zoomed images represent 50μm. Quantification of **(F)** cell numbers and **(G)** cell sizes for *Smedwi+* and gamma-neoblast in RNAi versus control animals at 5 days post RNAi. *** p<0.001 (2-way ANOVA).

**Fig. S3.**
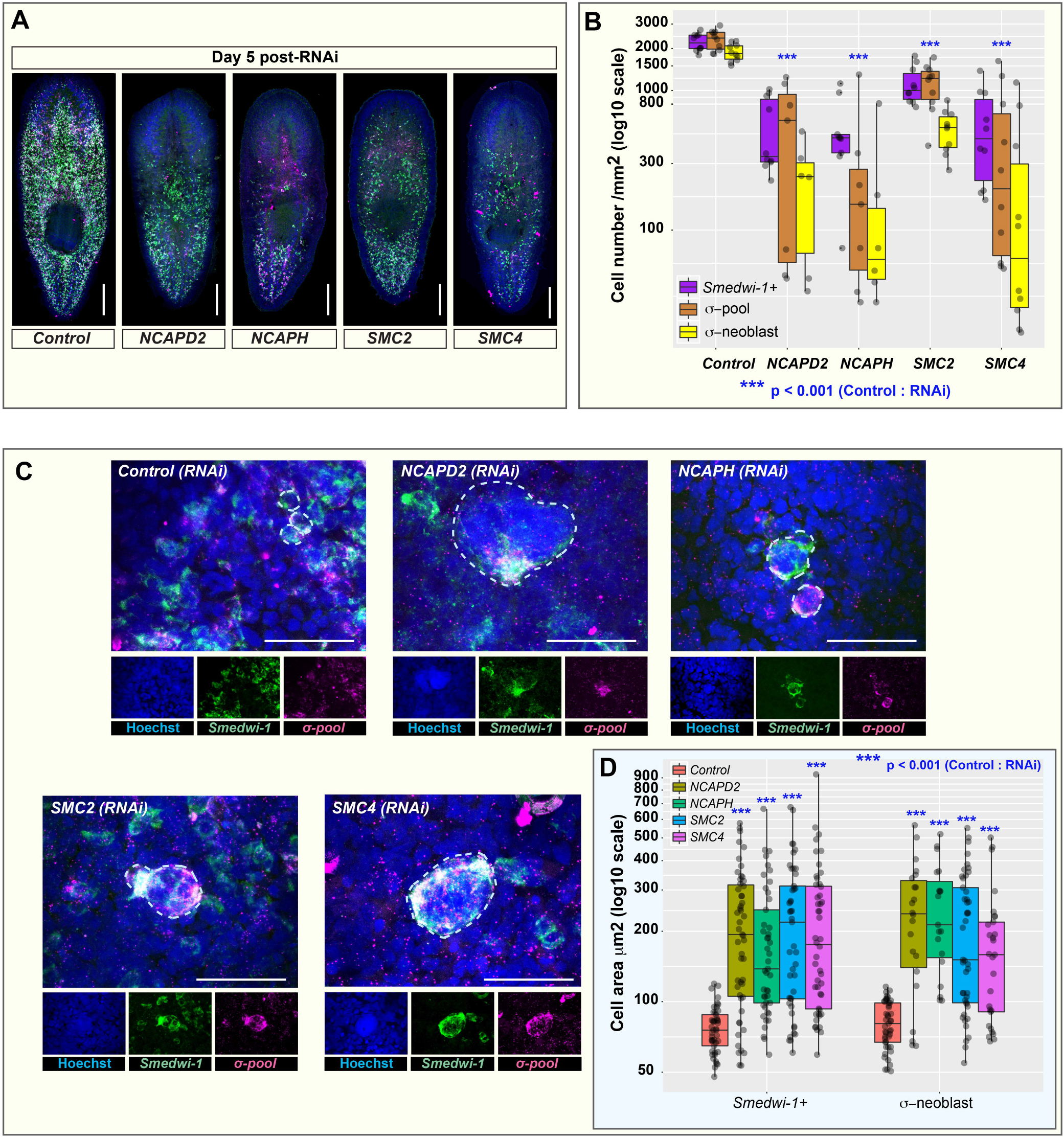
RNAi of condensin I subunit genes results in the enlargement and depletion of sigma-neoblasts. **(A)** Lineage markers labelling sigma-neoblasts were assessed by dFISH in *NCAPD2, NCAPH, SMC2* and *SMC4* RNAi animals at 5 days post RNAi. Scale bars represent 0.5mm. **(B)** Quantification of sigma-neoblasts in RNAi animals at the same time point. ***p < 0.001 (Student’s t-test; pairwise comparisons between each cell population in control versus RNAi animals). **(C)** A subset of sigma-neoblasts are enlarged and multinucleated in condensin RNAi animals. Scale bars represent 50μm. **(D)** Quantification of cell sizes for *Smedwi+* and sigma-neoblasts in condensin RNAi animals versus control animals at 5 days post RNAi. *** p<0.001 (Student’s t-test).

**Table S1**. List of primers used in PCR and cloning.

**Dataset S1**. Fasta file containing nucleotide sequences of condensin complex I subunits in triclad planarians.

